# Initial Characterization of the Two ClpP Paralogs of *Chlamydia trachomatis* Suggests Unique Functionality for Each

**DOI:** 10.1101/379487

**Authors:** Nicholas A. Wood, Krystal Chung, Amanda Blocker, Nathalia Rodrigues de Almeida, Martin Conda-Sheridan, Derek J. Fisher, Scot P. Ouellette

**Affiliations:** Department of Pathology and Microbiology, College of Medicine, University of Nebraska Medical Center, Omaha, NE; Division of Basic Biomedical Sciences, Sanford School of Medicine, University of South Dakota, Vermillion, SD; Department of Microbiology, Southern Illinois University Carbondale, Carbondale, IL; Department of Pharmaceutical Sciences, College of Pharmacy, University of Nebraska Medical Center, Omaha, NE

**Keywords:** *Chlamydia*, differentiation, protein turnover, protein quality control, Clp protease

## Abstract

*Chlamydia* is an obligate intracellular bacterium that differentiates between two distinct functional and morphological forms during its developmental cycle: elementary bodies (EBs) and reticulate bodies (RBs). EBs are non-dividing, small electron dense forms that infect host cells. RBs are larger, non-infectious replicative forms that develop within a membrane-bound vesicle, termed an inclusion. Given the unique properties of each developmental form of this bacterium, we hypothesized that the Clp protease system plays an integral role in proteomic turnover by degrading specific proteins from one developmental form or the other. *Chlamydia* has five uncharacterized *clp* genes: *clpX*, *clpC*, two *clpP* paralogs, and *clpB*. In other bacteria, ClpC and ClpX are ATPases that unfold and feed proteins into the ClpP protease to be degraded, and ClpB is a deaggregase. Here, we focused on characterizing the ClpP paralogs. Transcriptional analyses and immunoblotting determined these genes are expressed mid-cycle. Bioinformatic analyses of these proteins identified key residues important for activity. Over-expression of inactive *clpP* mutants in *Chlamydia* suggested independent function of each ClpP paralog. To further probe these differences, we determined interactions between the ClpP proteins using bacterial two-hybrid assays and native gel analysis of recombinant proteins. Homotypic interactions of the ClpP proteins, but not heterotypic interactions between the ClpP paralogs, were detected. Interestingly, ClpP2, but not ClpP1, protease activity was detected *in vitro*. This activity was stimulated by antibiotics known to activate ClpP, which also blocked chlamydial growth. Our data suggest the chlamydial ClpP paralogs likely serve distinct and critical roles in this important pathogen.

**Importance:** *Chlamydia trachomatis* is the leading cause of preventable infectious blindness and of bacterial sexually transmitted infections worldwide. Chlamydiae are developmentally regulated, obligate intracellular pathogens that alternate between two functional and morphologic forms with distinct repertoires of proteins. We hypothesize that protein degradation is a critical aspect to the developmental cycle. A key system involved in protein turnover in bacteria is the Clp protease system. Here, we characterized the two chlamydial ClpP paralogs by examining their expression in *Chlamydia*, their ability to oligomerize, and their proteolytic activity. This work will help understand the evolutionarily diverse Clp proteases in the context of intracellular organisms, which may aid in the study of other clinically relevant intracellular bacteria.

## Introduction

*Chlamydia trachomatis* is one of the most prevalent human sexually transmitted infections and the leading cause of preventable infectious blindness worldwide (1). Of particular note are the negative effects associated with untreated *C. trachomatis* infections. Because of the asymptomatic nature of 60-80% of cases (2), infection by this organism can lead to complications such as pelvic inflammatory disease, which can in turn lead to infertility and ectopic pregnancies in women (3). An infant can also acquire conjunctivitis during birth should the mother be infected, which can result in irreversible blindness if left untreated.

*C. trachomatis* is an obligate intracellular pathogen with a complex developmental cycle (see (4) for detailed review). These pathogenic bacteria differentiate between two distinct functional and morphological forms over the course of 48 to 72 hours, depending on the strain. The electron dense elementary body (EB) is the infectious but non-dividing form that is characterized by its histone-compacted chromosome (5) and cysteine-rich, disulfide-crosslinked outer membrane (6). The reticulate body (RB) is the non-infectious and replicating form that has a relaxed chromosome structure and lipid-based cell walls that are sensitive to disruption. The typical size of each form is ~0.3 μm and ~1 μm, respectively (4). A single EB initiates infection of a host cell by attaching to the plasma membrane and inducing uptake into the cell by type III secreted effectors (7) (8). The EB remains within a host-derived vesicle, which is diverted from the endocytic pathway, and differentiates into the much larger RB. The vesicle is rapidly modified into a pathogen-specified parasitic organelle termed an inclusion (9) (10). Proliferation of this RB within the inclusion follows until such time that secondary differentiation occurs, and the RBs begin to condense their genome and crosslink their outer membrane. After a significant amount of EBs have accumulated, they exit the cell for infection of surrounding cells. Proteomic and transcriptional analyses indicate that EBs and RBs have distinct patterns of gene expression and protein repertoires (11) (12) (13) (14). Given the striking phenotypic differences between the two developmental forms, we hypothesize that protein turnover plays a key role in differentiation in addition to general maintenance of bacterial homeostasis during both normal and persistent growth modes (15) (16) (17).

Even though *Chlamydiae* exhibit such a complex developmental cycle and evade the host immune system during chronic infections, these bacteria have undergone significant reductive evolution to eliminate unnecessary genes. This in turn suggests that most genes retained by this organism likely serve an important function to bacterial fitness (18). Within its genome, *Chlamydia* encodes homologs to four **<u>c</u>**aseino**<u>l</u>**ytic **<u>p</u>**rotease *clp* genes: *clpC*, *clpX*, *clpP* (*clpP1* and *clpP2*), and *clpB*. ClpB is a putative deaggregase and does not appear to interact directly with other Clps for proteomic turnover (12). ClpC and ClpX proteins are classified as AAA+ (**<u>A</u>**TPases **<u>A</u>**ssociated with various cellular **<u>A</u>**ctivities) unfoldases that serve as adaptor proteins to linearize target proteins in an ATP-dependent manner (19) (20). ATP binding, but not necessarily hydrolysis, potentiates interaction between a homo-hexamer of these proteins and a ClpP complex in other organisms (21) (22). ClpP is a serine protease that gives peptidase function to the resultant complex (23). ClpP proteins have been shown to oligomerize into a tetradecameric complex of a stack of two heptamers that can then perform proteolytic function upon interaction with the adaptor protein oligomer (24) (25). Binding of the unfoldase protein complexes relaxes and stabilizes the N-terminal region of the ClpP complex, allowing larger substrates into the proteolytic complex.

Multiple pathogenic bacteria possess two *clpP* paralogs including *Pseudomonas aeruginosa*, *Listeria monocytogenes*, and *Mycobacterium tuberculosis*. The Baker lab characterized the dual ClpP system of *P. aeruginosa* and demonstrated that each system potentially has a distinct function contributing to virulence and fitness of the bacterium (26). While the group studying ClpP1/2 in *Mycobacterium* showed heterologous interactions between these paralogs, these genes are co-transcribed, suggesting coupled function (27). The dual ClpP proteases of *Listeria monocytogenes* (Lm) were shown to form both hetero- and homotypic complexes (28) (29), yet the authors demonstrated that Lm ClpP2 likely can function independently of Lm ClpP1. These studies, taken together with the reductive evolution of the chlamydial genome and the distinct protein repertoires of EBs and RBs, led us to hypothesize that the chlamydial ClpPs may serve distinct and critical functions in the physiology of the organism.

To begin investigation of the role of the Clp protease system in chlamydial biology, we initiated a series of studies to characterize the ClpP paralogs. All *clp* genes are expressed as RB-specific gene products during the developmental cycle. Bioinformatic and structural modeling analyses indicate that ClpP1 and ClpP2 proteins retain key residues important for function as well as structure. Interestingly, over-expression of a mutant ClpP1 was detrimental to chlamydial development whereas a mutant ClpP2 had no observable effect. Over-expression of wild-type ClpPs had no effect on recovery of infectious EBs. To explore the basis of these observations, we performed a series of *in vitro* and *in vivo* assays to determine oligomerization and protease activity of each paralog. Whereas we observed homo-oligomerization of each ClpP paralog, we did not detect hetero-oligomerization between these proteins. From *in vitro* protease activity assays, we observed proteolytic activity for ClpP2 only. Consistent with this, antibiotics known to activate ClpP proteases stimulated the protease activity of ClpP2 only and blocked chlamydial growth. Combined, our data suggest ClpP1 activity may be tightly regulated. We conclude that each chlamydial ClpP protease serves a unique and independent role, either in differentiation or more conserved physiological processes.

## Results

### The *clp* genes are expressed as RB specific genes

We hypothesize, based on the arrangement of the genes in the chromosome, that ClpC interacts with ClpP1 and that ClpX interacts with ClpP2. This does not preclude each individual component from acting independently and is not mutually exclusive to our hypothesis. We base our prediction of ClpP2X interaction due to the juxtaposition of the two genes within the same operon (Fig. 1A). We then reasoned that ClpP1 would serve as the proteolytic subunit that would interact with ClpC, as both are encoded independently of each other in separate genomic contexts (Fig. 1A). Because chlamydiae are developmentally regulated bacteria, gene expression can differ from one developmental form to the other. Gene expression in *Chlamydia* can be broadly categorized into three different stages: early-, mid-, and late-developmental cycle genes (30). Early cycle genes are involved in initial differentiation from EB to RB and establishment of the nascent inclusion. Mid-cycle genes are RB specific and typically play a role in growth and division of the RBs as well as in inclusion modification. Late cycle genes function in differentiation from an RB to an EB or serve an important function in the secondary infection of a host cell. The specificity of a transcript to a particular point of the developmental cycle can give insight into the possible function of the encoded protein. To determine when during the developmental cycle the *clp* genes are transcribed, we measured the transcription of each gene over a time course of infection. Nucleic acid samples were collected for analysis at various time points of infection using wild-type *C. trachomatis* L2. Our data indicate that the *clp* genes analyzed are transcribed in a pattern consistent with mid-cycle, RB specific function since the transcript levels peak after primary differentiation has occurred and a population of RBs has been established (i.e. 16 hpi; Fig. 1B-E).

**Figure 1:**
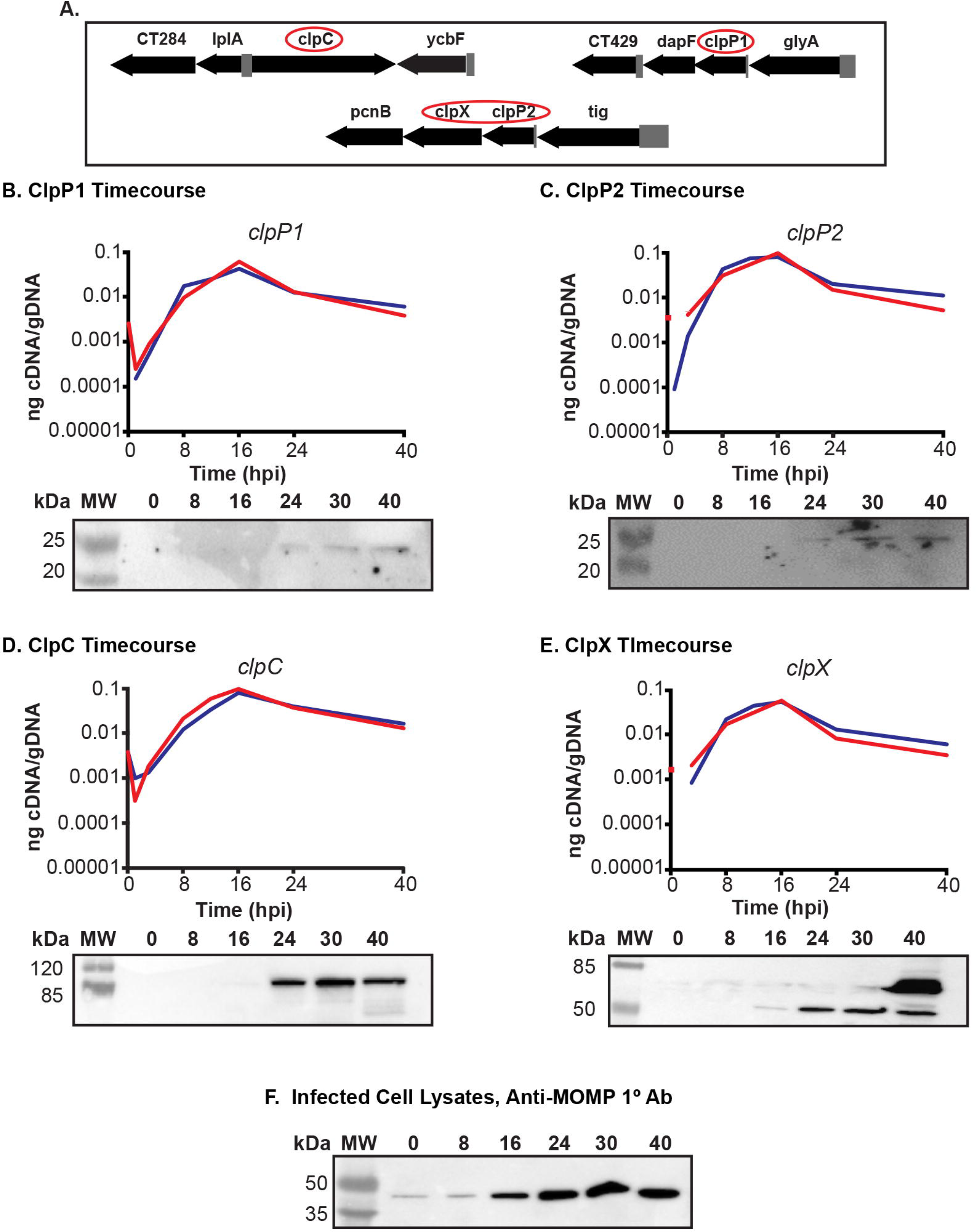
The *clp* genes are expressed during RB growth. **(A) Gene maps** of the *clp* genes in the *Chlamydia trachomatis* genome. Gene numbers are in numerical order from left to right and reflect the serovar D numbering scheme of Stephens et al. (18). The *clp* genes are circled in red. The *clpP2* gene is *ct706*. **(B) – (E) Temporal expression** of the *clp* genes. RT-qPCR analysis of the *clp* genes from two independent time course experiments of a *C. trachomatis* serovar L2 infection of HEp2 cells was performed. Total RNA and DNA were collected at the indicated times post infection and processed as described in Materials and Methods. Equivalent amounts of cDNA were used for each assay and analyzed in triplicate. Results were reported as a ratio of cDNA to genomic DNA. Standard deviations for each were typically less than 10% of the sample. Note that some transcripts were not detectable at 1 hpi. Western blotting was performed on whole cell lysates of total protein from a time course of *Chlamydia*-infected cells, separated by SDS-PAGE, and transferred to a nitrocellulose membrane for blotting. All four of the genes analyzed appear to be expressed mid-developmental cycle. **(F) Major outer** membrane protein (MOMP) was blotted as a control for chlamydial development over the course of infection.

In parallel, we analyzed lysates from infected cells using primary antibodies generated against the Clp proteins. Given the high level of homology between the *Pseudomonas aeruginosa* (Pa) ClpP1 and *C. trachomatis* (Ctr) ClpP2 (see Fig. 2), we speculated a polyclonal antibody developed against Pa_ClpP1 (a kind gift of Dr. T. Baker, MIT) would detect Ctr ClpP2 by western blot. As seen, Ctr ClpP2 was detected from infected cell lysates starting at 24h post-infection (hpi) using the polyclonal anti-Pa_ClpP1 antibody (Fig. 1C). Given the relatively low levels of ClpP2 compared to the major outer membrane protein (MOMP; Fig. 1F), we cannot conclude that ClpP2 is not present at earlier times or that an antibody specific to chlamydial ClpP2 would be more sensitive to detect such lower levels. The polyclonal antibody developed against Pa_ClpP2 did not react with chlamydial lysates or recombinant Ctr ClpP1 or ClpP2 (Fig. S1). We obtained additional antibodies raised against chlamydial ClpP1, ClpX, and ClpC (a kind gift of Dr. G. Zhong, UTHSC) and used these antibodies to detect their targets (validated in Fig. 1S). We observed similar patterns of expression for these proteins as for ClpP2 although faint bands for ClpX and ClpC were observed at 16hpi (Fig. 1B, D, & E). Clp protein levels mirror the transcripts and are detected from mid-developmental cycle for the remainder of the infection. These patterns are distinct to those of canonical early or late cycle genes such as *euo* or *omcB* (31) (32) (33). Operon status seems likely for *clpP2* and *clpX* given that *clpX* so closely mirrors *clpP2* with slightly lower transcript abundance. Overall, the expression data indicate that the Clp components are expressed primarily in the RB phase of growth and division. This does not, however, preclude these proteins from having functions at other times during the developmental cycle.

**Figure 2:**
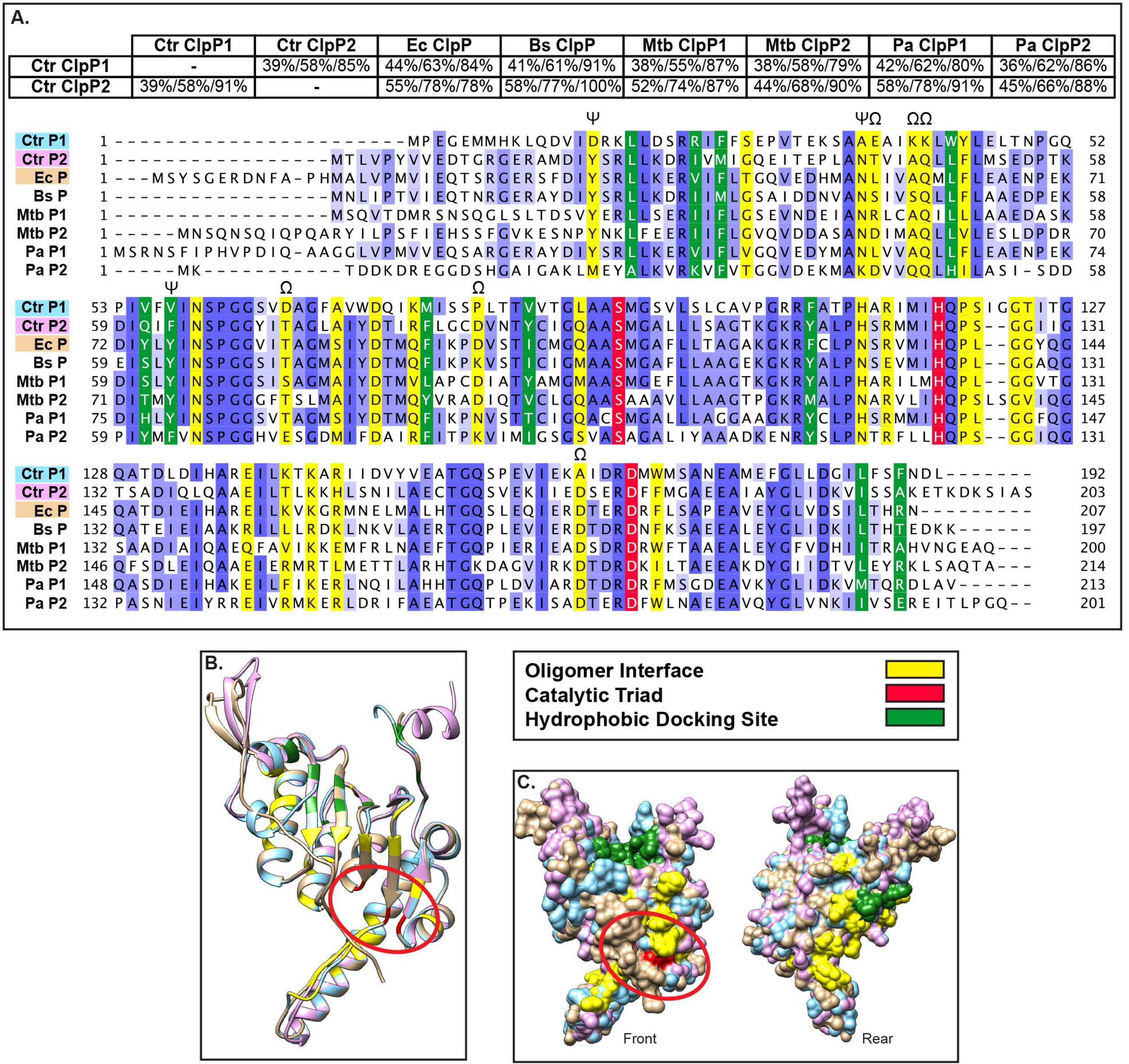
Bioinformatics analyses suggest both chlamydial ClpP paralogs are functional proteases. Pairwise alignments performed using NCBI-BLAST (default settings) and presented as %Identity/%Similarity/%Coverage. Multiple sequence alignment performed using Clustal Omega default settings and presented using Jalview Version 2. Organisms included are *C. trachomatis* (Ctr), *E. coli* (Ec), *B. subtilis* (Bs), *M. tuberculosis* (Mtb), and *P. aeruginosa* (Pa). Predicted 3D structures acquired from Phyre2 aligned and colored in Chimera (UCSF). **(A) Conserved residues** highlighted in varying shades of blue depending on conservation strength across species, with the catalytic triad in red, the oligomer interface in yellow, and the hydrophobic docking region in green. Residues in which ClpP1 has a radically different substitution compared to other ClpP proteins are denoted by Ω. Residues involved in activation conformational changes denoted by ψ (35) **(B) 3D structural alignment** of Ctr ClpP1, Ctr ClpP2, and Ec ClpP (peptide colors as shown in multiple sequence alignment). Active site residues are circled in red. **(C) Space filling model** of predicted 3D structures. “Rear” is a 180° rotation around the Y axis to show the surface of the protein not shown in the “Front” image.

### Bioinformatics analyses identify key residues important for structure and function of each chlamydial ClpP paralog

We hypothesized that each ClpP paralog of *C. trachomatis* serves distinct functions in the biology and pathogenesis of this unique bacterium, despite their similarity in expression patterns. To further test this hypothesis, we performed a series of bioinformatics analyses. We began by looking at pairwise alignments of the chlamydial ClpP proteins against each other and against ClpP homologs from other bacteria (Fig. 2A). Interestingly, each chlamydial ClpP shared more homology to ClpP of *E. coli*, at 44% and 55% identity with expect values of 4×10^−51^ and 2× 10^−83^, respectively, and to other ClpP homologs than to its paralog. This observation was supported by the ability of the anti-Pa_ClpP1 antibody to recognize Ctr ClpP2 but not Ctr ClpP1 (Fig. 1C). The sequence identity between Ctr ClpP1 and Ctr ClpP2 is 39% with an expect value of 10^−44^. Similarly, the two ClpP paralogs of *P. aeruginosa* also share a low amount of similarity to each other compared to other ClpP homologs via protein alignment (data not shown), and each serves a distinct function in *P. aeruginosa* (26). Therefore, these observations support the likelihood that the chlamydial ClpP paralogs have distinct roles.

A closer analysis of the chlamydial ClpP paralogs provides further evidence for unique functionality of each protein. A 3-dimensional predicted structure alignment of Ctr ClpP1 and ClpP2 to *E. coli* ClpP indicates that the structural architecture would allow for homo-oligomer formation and interaction with unfoldases (Fig. 2B), though distinct differences in the oligomer interface may prevent hetero-oligomerization (34). Additional variation in structure is mostly limited to the N- and C-termini, however the catalytic triad active site aligns well for all ClpP homologs analyzed (Fig. 2B&C). Also unique to chlamydial ClpP1 are distinct differences at residues that align to amino acids critical to activation in other bacteria. An alanine at position 37 of Ctr ClpP1 may result in a conformational difference in the P1 pre-activation complex that precludes P1/P2 interaction (35), which may act as a layer of regulation. Further supporting differential regulation is the lack of an aromatic amino acid at position 57 of ClpP1 that is relatively well-conserved in other ClpP homologs, suggesting that Ctr ClpP1 enters the “open” conformation differently than Ctr ClpP2. Given the conserved catalytic residues and the structural similarity of the Ctr ClpP proteins to homologs from a diverse set of bacteria, we conclude that the chlamydial ClpP proteins are *bona fide* proteases. However, the predicted structural differences at the termini of the protein and in other key residues suggests that the chlamydial ClpP proteins (i) do not interact with each other and (ii) have specialized functions.

### Overexpression of only inactive ClpP1 negatively affects *C. trachomatis*

Leveraging recent advances in chlamydial genetics, we next determined the effect of over-expression of ClpP proteins in *C. trachomatis*. We designed and constructed plasmids encoding wild-type or inactive mutants of each *clpP* paralog, fused to a 6xHis tag, under the control of an anhydrotetracycline (aTc)-inducible promoter (36). We successfully transformed *C. trachomatis* with plasmids encoding either ClpP1 S92A or ClpP2 S98A active site serine mutants that are known to abolish protease activity (See Fig. 2B; (37)) in addition to plasmids encoding either wild type ClpP protein. To assess the effect of over-expression of both the wild type and the inactive mutant ClpPs, HEp2 cells were infected with each transformant, and expression was induced at 10hpi with 10 nM aTc. 14-hour pulses were utilized to determine inclusion morphology at 24hpi. Use of aTc at 10 nM did not alter wild-type chlamydial inclusion growth (Suppl. Fig. S2; (36)). At that time point, the localization of the ClpP proteins was within the bacterial cytosol. Over-expression of mutant ClpP1 resulted in noticeably smaller inclusion sizes after 14h of induction (Fig. 3A) whereas over-expression of inactive ClpP2 had no demonstrable effect (Fig. 3B). Of note is that neither of the wild type chlamydial ClpP proteins had any apparent negative impact. Further analysis of recoverable inclusion forming units (IFUs) confirmed the observed reduction in chlamydial viability following ClpP1 S92A overexpression, with a severe decrease in IFU recovery following a 14h pulse with the aTc inducer (Fig. 3C). Conversely, ClpP2 S98A or wild-type protein overexpression has no statistical and likely no biologically significant impact on development. From these data indicate we conclude that over-expression of ClpP1 S92A is detrimental to chlamydiae. In support of this, we were unable to isolate a clonal population of purified ClpP1 S92A transformants as we routinely observed small bacterial populations susceptible to antibiotic selection even after multiple passages. This observation indicates that leaky expression may drive plasmid loss as has been observed in other chlamydial transformation systems (38) (39).

**Figure 3:**
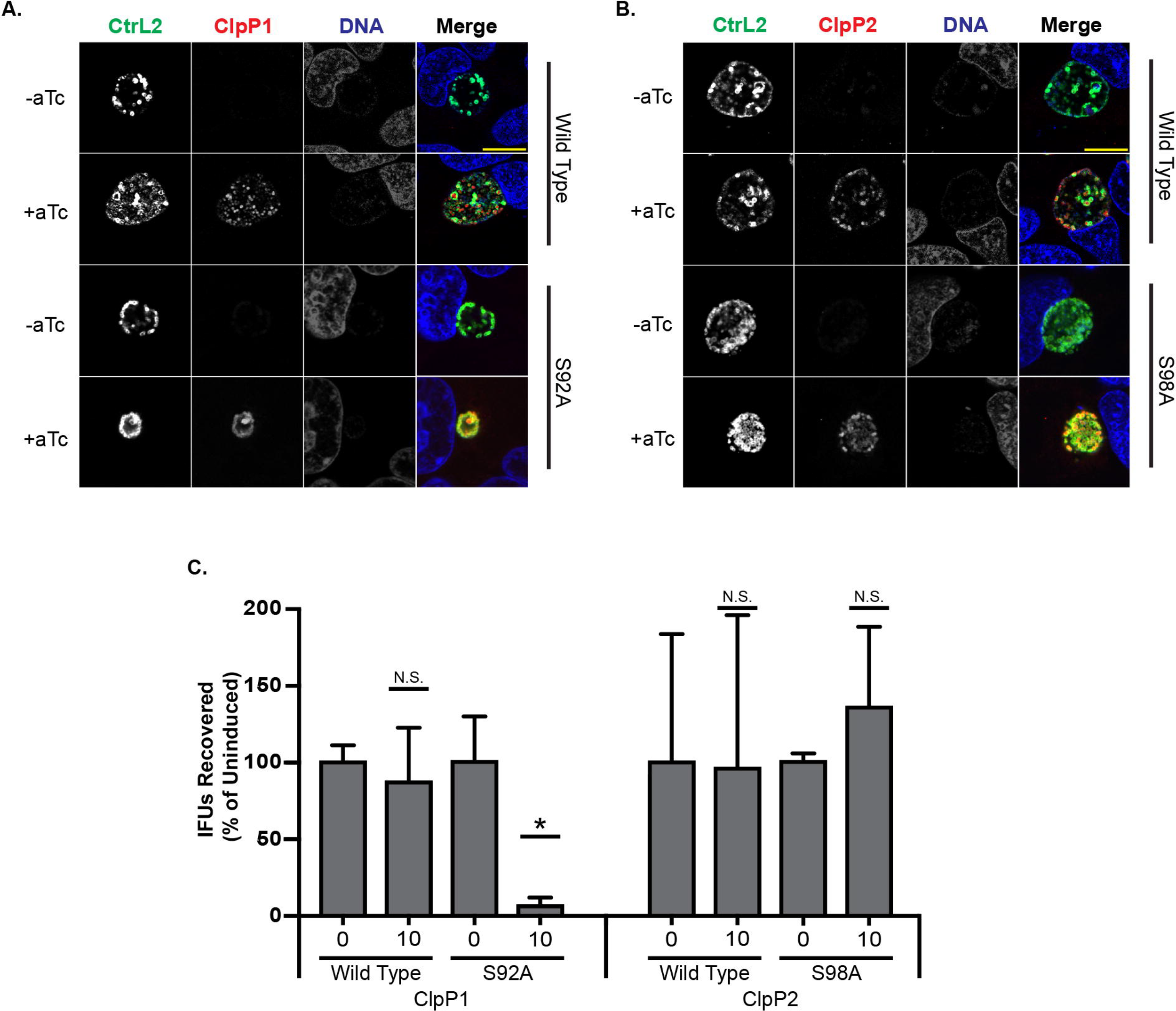
Overexpression of wild type or inactive ClpP proteins has varying effects on *Chlamydia*. *C. trachomatis* serovar L2 was transformed with anhydrotetracycline (aTc)-inducible shuttle vectors encoding either wild type or active site mutants of each ClpP paralog with a 6xHis tag at the C-terminus. HEp2 cells were infected with each transformant, and expression was induced at 10h post infection (hpi). **(A) ClpP1 wild type and S92A overexpression** assay. Overexpression of inactive ClpP1 has a negative impact on the bacteria. **(B) ClpP2 wild type and S98A overexpression** assay. Parameters same as described above. Overexpression does not appear to negatively affect *Chlamydia*. Samples were stained for major outer membrane protein (MOMP; green), 6xHis tagged ClpP protein of interest (red), and DNA (blue). Representative images of three independent experiments are presented. Scale bars are equal to 10 μm. Images were acquired on a Zeiss LSM 800 laser scanning confocal microscope with a 60x objective and a 3x digital magnification. **(C) Inclusion forming unit (IFU) assay** measuring the effect of increasing levels of ClpP protein induction on chlamydial growth. Values and error bars are averages of three independent experiments and are reported as a % of the respective uninduced sample. ^*^= P<0.05, N.S.= not significant.

### The ClpP paralogs demonstrate homotypic, but not heterotypic, interactions

To better understand the function of each ClpP paralog and why over-expression of ClpP1 S92A, but not ClpP2 S98A or wild-type proteins, was detrimental to *Chlamydia*, we initiated a series of *in vivo* and *in vitro* biochemical assays to characterize the properties of the ClpP proteins. Extensive evidence exists from other bacterial systems to indicate that ClpP forms a tetradecameric complex composed of two heptameric rings (21-25, 40, 41). To determine whether the chlamydial ClpP paralogs formed homo- and/or heterooligomers, we first employed the bacterial adenylate cyclase-based two-hybrid (BACTH) system to detect protein-protein interactions. This *in vivo* assay is based on the reconstitution of adenylate cyclase activity when two functional fragments of Cya from *B. pertussis* are brought into close proximity by interacting proteins (42). Interactions can be qualitatively determined on Xgal plates and quantified by beta-galactosidase activity. Using the BACTH system, we observed homo-oligomerization of each ClpP paralog, but failed to detect interactions between ClpP1 and ClpP2 (Fig. 4A&B). Importantly, the active site mutants also interacted with each other and the wild-type proteins, indicating that this mutation did not interfere with homo-oligomerization (Fig. 4B). These data are consistent with the predicted structural and sequence differences noted (Fig. 2B&C).

**Figure 4:**
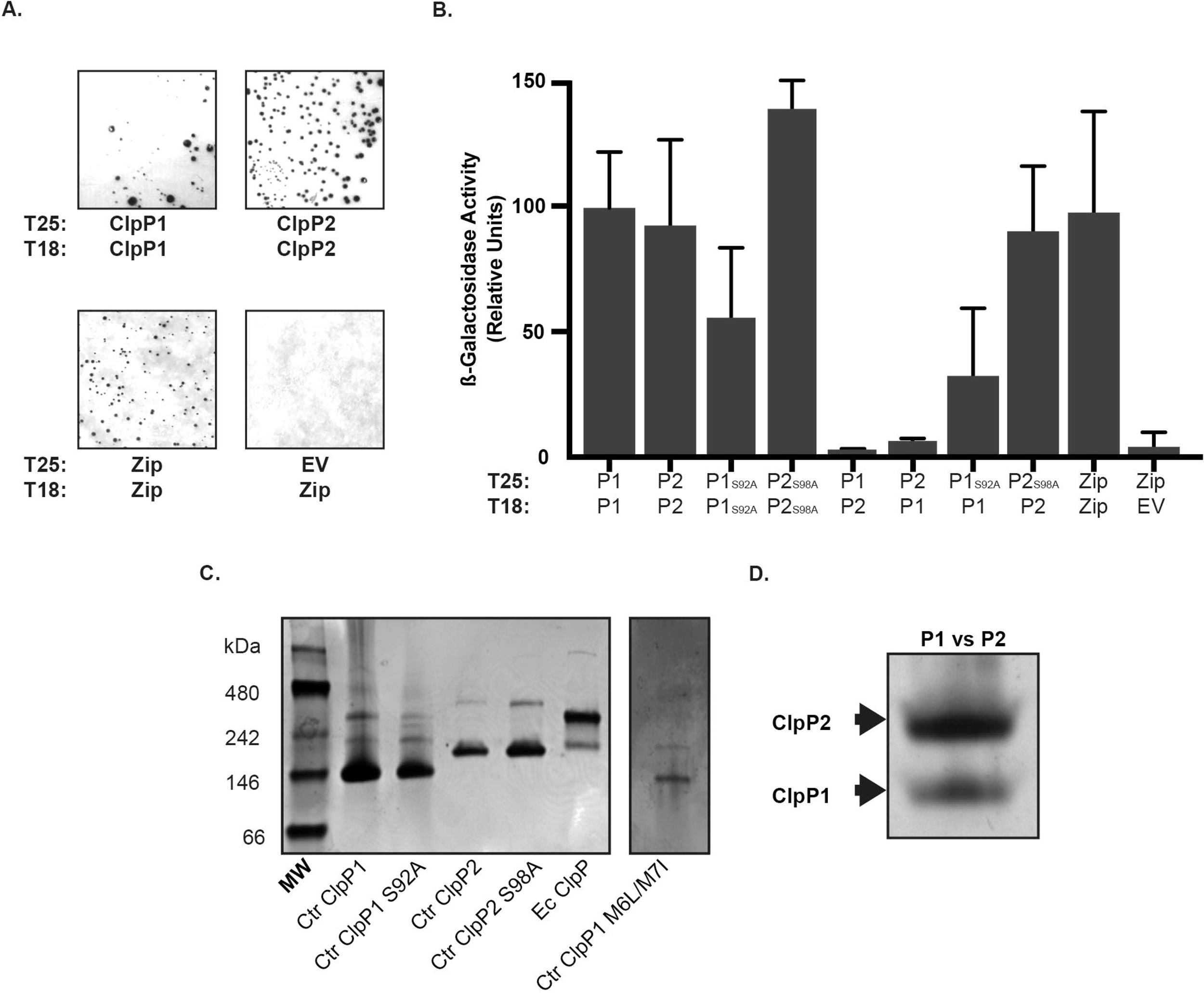
The chlamydial ClpP paralogs self-associate but do not form hetero-oligomeric complexes. **(A) Bacterial adenylate cyclase-based two-hybrid assay** of homotypic interactions of *C. trachomatis* (Ctr) ClpP1 and ClpP2. The indicated constructs encoding each *clpP* paralog fused to the T18 or T25 fragment of the Cya toxin of *B. pertussis* were co-transformed into Δ*cya E. coli* DHT1 and plated on minimal medium containing X-gal. Interactions between proteins results in the formation of blue colonies. Representative images of results from at least three independent experiments are shown along with positive (Zip/Zip) and negative (T25 empty vector versus T18-Zip) controls. Photos set to grayscale. **(B)** β**-Galactosidase assay** results from experiments performed as described in Panel (A). Y-axis is a measurement of relative units of beta-galactosidase activity. X-axis indicates the test conditions for proteins fused to the T25 or T18 as indicated. Interactions are considered positive when at least five times the activity of the negative control is measured. Only homotypic interactions were positive in these assays. **(C) Test of ClpP Oligomerization by Native-PAGE**. 5 μg samples were run on 4-20% native-PAGE gels and stained with Coomassie for protein detection. Representative results from at least three experiments with independent protein purifications are shown. The Ctr ClpP1 methionine mutant (M6L/M7I) was run in a separate lane and the contrast was uniformly adjusted to help with band visualization. Native molecular weight markers are shown to the left of the gel. *E. coli* (Ec) ClpP is included as a positive control on the far right of the gel. **(D) Test of Hetero-oligomerization between Ctr ClpP1 and Ctr ClpP2.** Each recombinant protein was incubated together prior to electrophoresis. The gel has been cropped and enlarged to aid in detecting P1 and P2 hetero-oligomers, which appear to be absent consistent with panel (B) results. The predicted heptamer/tetradecamer sizes in kDa are: Ctr ClpP1, 154/308; Ctr ClpP2 161/322; and Ec ClpP 168/336.

As the BACTH system cannot indicate oligomerization state, we next purified recombinant chlamydial ClpP1 and ClpP2 (22 and 23 kDa, respectively), including the active site mutants, and analyzed their migration by native PAGE. As a positive control, we also purified *E. coli* ClpP (24 kDa). Each of the recombinant chlamydial proteins clearly formed a heptamer with a fainter, slower migrating band indicating potential tetradecamer forms (Fig. 4C). The *E. coli* ClpP, in contrast, migrated primarily as a tetradecamer with a less intense band at the predicted size of a heptamer. In purifying Ctr ClpP1, we noticed the presence of a doublet band (Suppl. Fig. S3). However, the N-terminus encodes two additional methionine residues at positions 6 and 7, thus the smaller product may have resulted from an alternative start site when expressed in *E. coli*. Mass spectrometry analysis of the N-terminus of each band supported this conclusion (Suppl. Fig. S3). Note that the BACTH constructs relied on N-terminal fusions such that the alternative start site of ClpP1 would not be relevant. To ensure that only the larger, full-length recombinant ClpP1 product was produced, we mutated the methionine residues to leucine and isoleucine. This recombinant ClpP1 (M6L/M7I) also migrated predominantly as a heptamer in the native PAGE assay (Fig. 4C). Therefore, we conclude that the smaller product did not alter the oligomerization state of the protein.

Both our bioinformatics predictions and the BACTH data indicated that ClpP1 and ClpP2 do not interact. To further test this, we mixed each recombinant protein in equimolar amounts and analyzed their oligomerization state by native PAGE. In support of our early observations, we did not observe heteromers of each ClpP paralog as each protein ran as distinct heptamers with no intermediate bands indicative of mixing of the monomers (Fig. 4D). However, we can neither exclude that the *in vitro* conditions preclude an interaction between these components nor that these paralogs interact *in vivo*. Nevertheless, given the differences in predicted structures at the termini of each ClpP paralog (Fig. 2) and the different interaction data presented, the parsimonious interpretation is a model wherein each protease complex functions separately.

### Chlamydial ClpP1 and ClpP2 vary in protease activity *in vitro*

To determine the functionality of the protease complexes, we used the recombinant ClpP proteins to assess their ability to degrade the small, fluorogenic compound Suc-LY-AMC. The degradation of this compound by other bacterial ClpP enzymes does not rely on ClpP activation (43), thus we could probe the basal protease activity of the ClpP paralogs. While ClpP2 degraded the substrate under our given assay conditions, ClpP1 activity could not be detected (Fig. 5A). This result for ClpP1 was not due to the presence of the alternative start site product since the M6L/M7I mutant displayed no activity in this assay. We also observed enhanced protease activity for ClpP2 in a sodium citrate buffer, as noted by others (44), but this buffer did not stimulate ClpP1 activity. That ClpP1 lacks the ability to degrade Suc-LY-AMC suggests the N-terminal region of the complex (Fig. 2) may block entry of even this small reporter substrate into the active site. Importantly, the activity of the ClpP2 protease was abolished by the active site serine mutation S98A (Fig. 5A). As a positive control, we also tested the activity of the *E. coli* ClpP complex in our assay conditions and observed degradation of the Suc-LY-AMP compound that was increased by sodium citrate (Fig. 5B). Thus, in spite of the activity of Ctr ClpP2, it displayed a significantly less proteolytic capacity *in vitro* compared to the *E. coli* ClpP. The apparent preference of heptameric formation by both chlamydial ClpP1 and ClpP2 could explain the reduced activity as compared to the *E. coli* ClpP, which preferentially formed tetradecamers *in vitro*. These observed differences may suggest that either the assay conditions are not optimal or that *Chlamydiae* tightly regulate the activity of their ClpP complexes.

**Figure 5:**
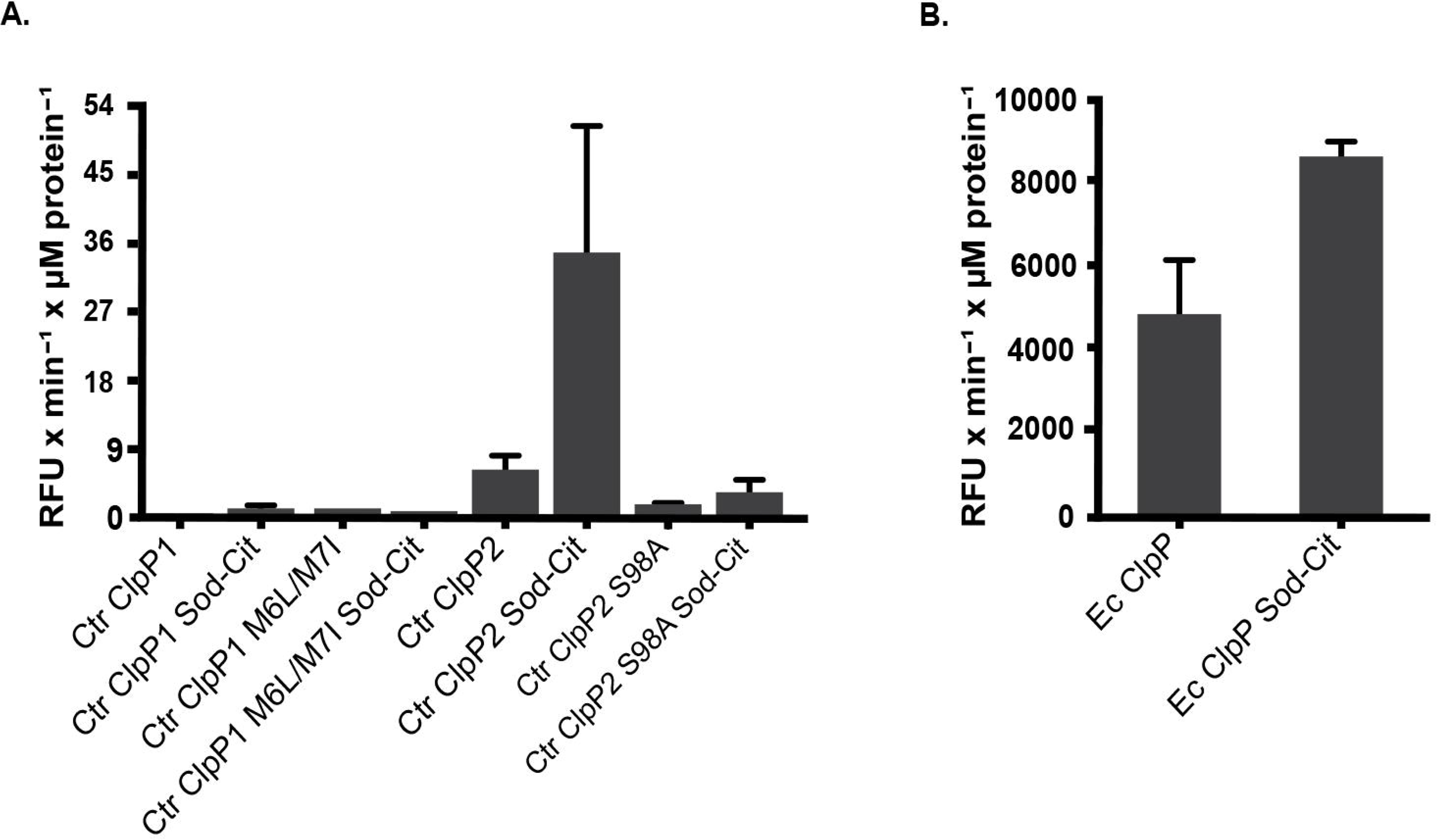
Chlamydial ClpP2, but not ClpP1, has ATPase-independent protease activity. *In vitro* protease activity of the *C. trachomatis* (Ctr) ClpP proteins versus a peptide substrate. ClpP samples (1 μM monomer) were incubated with the fluorometric peptide Suc-Luc-Tyr-AMC, and fluorescence was analyzed over a time. **(A) Relative activity** is shown for each protein run with or without the sodium citrate buffer. Use of sodium citrate buffer is indicated as “Sod-Cit”. Loss of activity is observed for the Ctr ClpP2 catalytic serine mutant. **(B) *E. coli* (Ec) ClpP** was used as a positive control. Experiments were run a minimum of three times and baseline assay values obtained with buffer alone were subtracted from each sample. Average values are reported with standard error.

### ClpP-activating antibiotics stimulate protease activity of ClpP2, but not ClpP1 and block chlamydial growth

Recently, a novel class of antibiotics that target ClpP have been developed (45). ACP and ADEP derivatives activate ClpP by eliminating or reducing its need for ATPase binding. The unregulated proteolysis that results has been demonstrated to eliminate persister bacteria (46). We used these compounds to determine whether the activated chlamydial ClpP proteases could degrade a more complex substrate, casein. As noted with the Suc-LY-AMP substrate, ClpP1 failed to degrade casein in the presence of any of the ACP1 derivatives (Fig. 6A). By contrast, the ACP1 compounds stimulated ClpP2 to degrade casein (Fig. 6B), albeit with some detectable substrate remaining at the end of the assay. As before, the *E. coli* ClpP was more efficient in degrading the substrate with little to no casein detectable (Fig. 6C; (47)). ClpP2 was thus able to, in the presence of these drugs that relax the N-termini of the complex, degrade larger substrates with intrinsically disordered regions (i.e. casein). The inability to detect proteolytic activity of ClpP1 again suggests that the N-terminus of ClpP1 may act as a regulatory block consistent with the differences we observed in our bioinformatics analysis (Fig. 2). Also consistent with the difference in activation is the lack of an aromatic amino acid at residue 57 of Ctr ClpP1. This particular residue of the hydrophobic pocket has recently been shown to be instrumental in ADEP-induced dysregulation of proteolytic activity, which supports a model where chlamydial ClpP1 is unaffected by activating drugs (35). However, we cannot exclude that we have not identified the optimal conditions necessary to demonstrate the enzyme activity of wild-type ClpP1.

**Figure 6:**
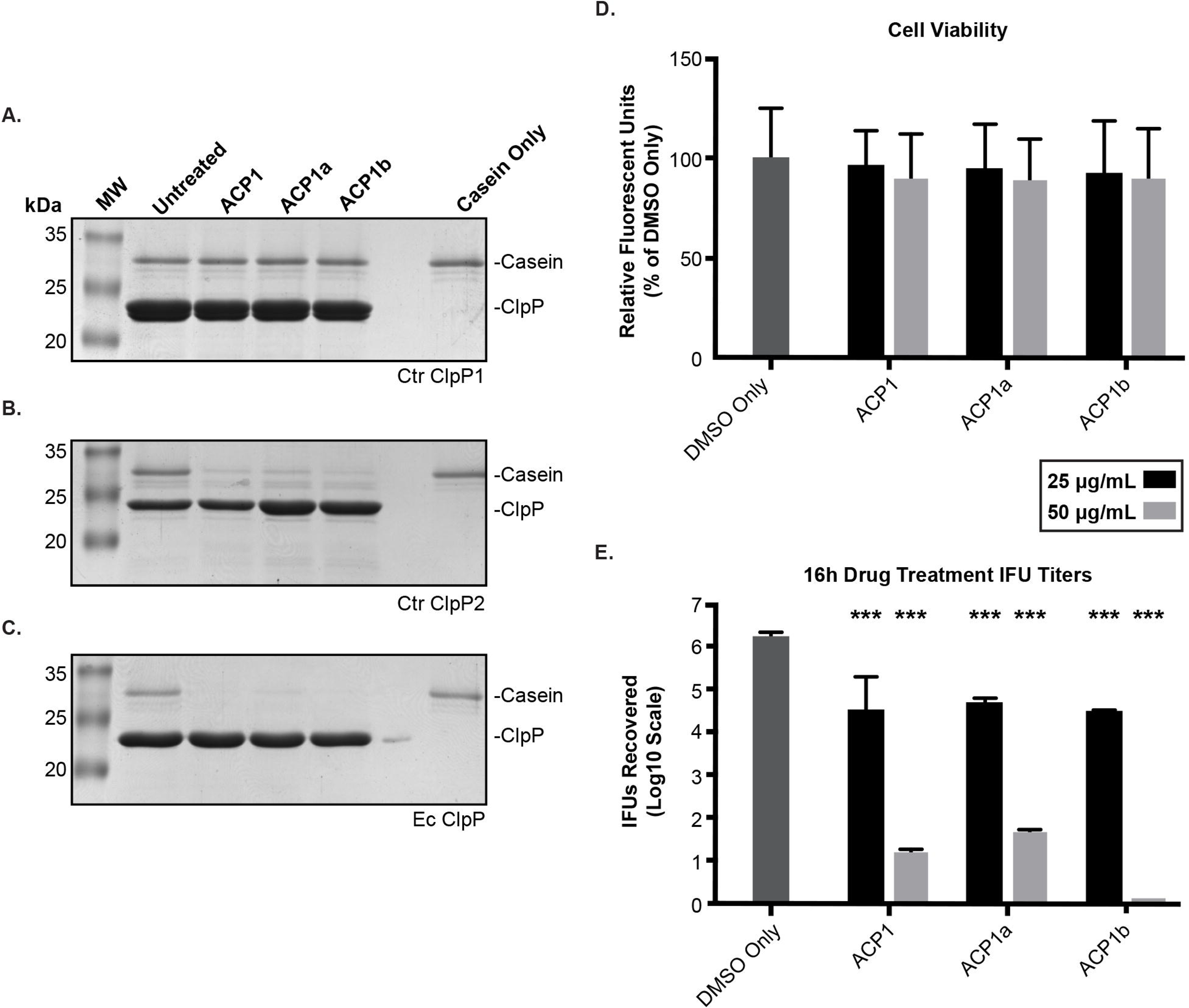
ClpP2, but not ClpP1, activity is stimulated by the antibiotic ACP1, which is detrimental to chlamydial growth. **(A-C) Casein (1 μg)** was incubated with 1 μM of the respective ClpP at 37 °C for 3 hours with or without the ACP compounds (used at 500 μM). Reactions were halted by mixing samples with Laemmli followed by boiling. Samples were run on 12% SDS-PAGE and stained with Coomassie. Molecular weight markers are shown to the left of each gel. Representative gels are shown; at least three experiments were performed for each protein. Ctr = *C. trachomatis.* Ec = *E. coli* **(D) Cell viability** of ACP treated cells were analyzed with resazurin assays. Values are reported as a percentage of the vehicle control and are representative of three independent experiments. **(E) Reinfection models** of ACP drug-treated cells reported on a Log_10_ scale. A student’s two-tailed T test was used to compare each parameter to the vehicle control (^***^= P<0.0001). For (D) and (E), HEp2 cells were infected with *C. trachomatis* serovar L2, and the ACP1 compounds or DMSO only was added to the infected cultures at 8 hours post-infection (hpi). At 24 hpi, cell viability was assessed for (D), or infected cells were collected to re-infect fresh monolayers to determine the recovery of inclusion forming units (IFUs) as described in Materials and Methods for (E).

As we were able to demonstrate the ability of the ACP1 compounds to activate Ctr ClpP2, we next sought to test the effect of these compounds on *Chlamydia.* To ensure that any effect on the bacteria was not due to host cell death, we first tested cell viability of infected, ACP-treated HEp2 cells. Indeed, we noted limited loss (~10%) in host cell viability for both 25 and 50 μg/mL concentrations following 16 hours of drug treatment (Fig. 6D), which reinforces that any loss of bacterial viability would not be due to a reduction in host cell viability. The ACP drugs all showed a remarkable impact on *Chlamydia* at both concentrations tested when added at 8hpi and samples were harvested at 24hpi (Fig. 6E). The lower drug concentration resulted in a 50-fold reduction in recoverable EBs whereas the higher concentration led to almost complete abrogation of chlamydial viability. Given the effects of the ACP compounds on ClpP protease activity *in vitro*, these data suggest that uncoupling of ClpP2 activity from its cognate ATPase strongly inhibits bacterial growth.

## Discussion

The Clp protease system has been extensively characterized in *E. coli* and *Bacillus subtilis*, and its importance in the growth and virulence of various pathogens (opportunistic or not) has been examined (see (23) for review). Since the discovery of the ClpP protease (48), interaction studies demonstrated that the ClpP proteins typically form a tetradecameric stack of two heptamers (49). The serine active sites are buried within these barrel-like structures (50). ClpP hydrolysis of large protein substrates is increasingly stimulated in the presence of AAA+ ATPase binding (51). However, complexed ClpP alone processes small substrates of up to 30 amino acids regardless of the presence of ATP, suggesting that active site availability and complex accessibility are the two main mechanisms regulating ClpP activity (52). Substrates that enter this complex by either ATPase-mediated or ATPase-independent means are hydrolyzed into short peptides of 6-8 amino acids with some degree of amino acid specificity (53) (44). While not unprecedented, the presence of two encoded *clpP* paralogs in a bacterial chromosome is unusual. In *P. aeruginosa*, another bacterium encoding dual ClpP proteases, the two isoforms have not been shown to interact in any test condition, and the different transcriptional profiles and differential effects on virulence factor production indicate these paralogs likely have distinct roles in *P. aeruginosa* (26). Conversely, the two ClpP proteins of *M. tuberculosis* and *L. monocytogenes* heterologously interact as two homotypic heptamers, which facilitates an increase in function (27, 29, 54). The activation of one homotypic ClpP heptamer activates the heptamer of the other isoform in *M. tuberculosis*, providing a novel mechanism for activation (41). Of note is that *L. monocytogenes* ClpP2 may also form a completely homotypic tetradecamer of one ClpP paralog (55). Again, the Clp proteolytic subunits play an integral role in virulence factor regulation in addition to enhancement of bacterial survival during intracellular growth (56). Taken together, these studies demonstrate the diverse function of dual ClpP peptidase systems in the pathogenesis and growth of various bacteria.

The work presented here to characterize the ClpP paralogs is the first to explore the function and role of the Clp protease system in *C. trachomatis*. Our results suggest that the two ClpP proteases likely serve distinct roles in chlamydial physiology and, we hypothesize, in the complex developmental cycle of this organism. We utilized bioinformatics techniques as an initial approach to investigate the potential function of the ClpP proteins. We observed significant similarity of the chlamydial ClpP paralogs to ClpP homologs of more studied organisms such as *E. coli*. In support of this, most of the hallmark conserved residues, including the active site triad, are present in the chlamydial ClpP paralogs. Both ClpP1 and ClpP2 have hydrophobic residues aligned with those of other bacterial ClpPs, suggesting the presence of AAA+ adaptor protein docking sites (57). Indeed, chlamydial ClpC and ClpX encode the evolutionarily retained IGF/L loop motifs that facilitate interaction with the ClpP tetradecameric complex (unpublished observation) (41). In spite of the similarities between the ClpP paralogs, there are also notable differences. The C-termini of both chlamydial ClpPs contain two residues that comprise the hydrophobic pocket (47, 58), but the alignment shows the presence of charged residues (D15, K40, K41, and D66) in ClpP1 while the conserved residues appear to be uncharged. In addition, ClpP1 lacks a highly conserved tyrosine residue (D15), a conserved glutamine (A36), and a conserved aromatic residue (V57), all of which contribute to canonical activation and tetradecamer chamber access for other homologs (35). Ongoing studies are investigating these residues in addition to others that are involved in complex activation. Of note is the predicted alpha helix at the N-terminus of ClpP1 and the C-terminus of ClpP2, which may contribute to some form of steric hindrance to prevent interaction between ClpP1 and ClpP2 (59). Indeed, we did not detect an interaction between ClpP1 and ClpP2 using both *in vivo* and *in vitro* techniques. Rather, we detected only homooligomers. While our *in vitro* assays have failed to show P1/P2 complex formation, we cannot rule out the possibility of protein modification promoting heteromers *in vivo*. Further studies examining the role of the N- and C-termini will be conducted to determine whether or not they play a role in complex formation and/or specificity.

Given the role of the N-terminal regions in regulation of the ClpP protease via reduced access into the proteolytic complex (60), the hypothesis of a difference in interaction or regulation of the ClpP complexes in *C. trachomatis* is reasonable based on the data presented here. We detected ClpP2 protease activity against both a small oligopeptide substrate and the more complex casein substrate. However, we did not detect proteolytic activity of ClpP1 *in vitro*, even in the presence of well-characterized drugs that activate ClpP complexes in the absence of an ATPase, which agrees with an important and highly regulated role for ClpP1 in *C. trachomatis*. The structural conservation and presence of the catalytic active site makes it unlikely that ClpP1 is a catalytically inactive protease. Rather, some form of modification or conformational change in structure, possibly chaperone or adaptor mediated, may be necessary to promote full complex formation and activity both *in vitro* and *in vivo*. In this model, an unknown regulatory factor may modulate ClpP1 activity. In a manner consistent with our bioinformatics data, ClpP1 may be activated non-canonically, as the mechanism needed to open the channel at the V57 residue is likely different than those targeting the aromatic amino acids present in other homologs. We are investigating this and other possibilities.

Our data lead us to hypothesize that each ClpP protein in *Chlamydia* forms a separate protease complex and, therefore, serves a different function. Whether each ClpP interacts specifically with ClpX or ClpC remains to be determined. However, the genomic proximity of ClpP2 to ClpX suggests a likely interaction with ClpX by sequestration immediately following translation; thus, we speculate that ClpP1 likely interacts preferentially with ClpC. Our qPCR and western blot data suggest that the *clpX*-encoding gene follows the *clpP2*-encoding gene in an operon, further supporting coupled function and an *in vivo* complex of ClpP2 and ClpX oligomers. Complex formation is currently under investigation as is what role each complex has *in vivo*. It is also possible that ClpP tetradecamers may function independently *in vivo* in a housekeeping role to degrade small oligopeptides resulting from degradation of larger proteins and/or import from the host cell. *Chlamydiae* encode a large repertoire of oligopeptide transporters, and the short peptide substrates are an important source of nutrients, particularly early during infection (61).

We were able to transform *C. trachomatis* with plasmids encoding either wild type or mutant ClpP1 or ClpP2. Overexpression of either wild type protein had seemingly little negative effect on growth and morphology, which strengthens a scenario where these proteins are tightly regulated and exhibit little function without a cognate ATPase. The tolerance of *C. trachomatis* to wild type ClpP1 and ClpP2 overexpression suggests that this system resists AAA+ ATPase saturation with an overabundance of either proteolytic subunit. Given the reduced tendency for either ClpP to form a tetradecamer, excess ClpP1 or ClpP2 beyond what is needed for homeostasis may also remain in a functionally inactive heptameric form unless recruited by an unfoldase. Interestingly, overexpression of the inactive ClpP mutants in *Chlamydia* resulted in differential effects on chlamydial growth and morphology. ClpP2(S98A) has, even upon the highest level of induction, little negative effect on inclusion morphology. Incorporation of inactive forms of ClpP2 (activity loss confirmed *in vitro*, Figure 5B) into the tetradecameric oligomer is not necessarily harmful to *C. trachomatis*. The ability of ClpP2 to degrade a small peptide reporter (Fig. 5A) is in congruence with a model where an unfoldase-independent ClpP2 tetradecamer targets and degrades imported peptides. Given that overexpression of inactive ClpP2 seems to have limited negative impact on chlamydial development, an inactive ClpP2 oligomer may bind small peptides, preventing extensive targeting of other, more vital substrates and negating any dominant negative effect. Conversely, overexpression of ClpP1(S92A) and its incorporation into the endogenous ClpP1 proteolytic machinery has a clear negative impact on *C. trachomatis*. Because overexpression of wild-type ClpP1 has no obvious negative effect, that inactive ClpP1 overexpression reduces chlamydial viability may mean that fully functional operation of ClpP1 is essential to the bacteria. The adverse effect observed may also suggest that ClpP1 has a limited reserve of unique adaptors that we have not yet identified and that production of an inactive ClpP1 complex titers out these adaptors and reduces function to the point of harm to the bacteria. We observed that the mutant ClpP proteins interacted with the wild-type proteins in the BACTH assay (Fig. 4B), suggesting the potential of the inactive proteins to form complexes with the endogenous proteins. Overall, these data suggest that, should each mutant protein oligomerize with endogenous wild-type protein *in vivo*, subunit poisoning of the ClpP1 complex is deleterious to chlamydiae whereas the ClpP2 complex is more tolerant.

We hypothesize that proteolytic Clp subunit regulation by a chaperone protein is vital to chlamydial survival. Based on our studies with the ACP1 antibiotics, our assays showed that ClpP2 acts in a manner similar to other well-characterized ClpP proteases when artificially activated by these compounds (Fig. 6A, C). Consequently, treatment of *Chlamydia*-infected cells with ACP1 (or one of its derivatives) significantly and drastically reduced chlamydial growth (Fig. 6E), which we speculate is due to the effects on ClpP2 function. Whether the negative effect stems from dysregulation of ClpP2 leading to uncontrolled proteolysis, which is supported by the *in vitro* degradation of casein, or from blocking of ClpP2/AAA+ ATPase complex formation, resulting in the inability to degrade larger substrates not accessible to ClpP2 alone, is currently under investigation. In spite of the inability of ClpP2(S98A) overexpression to significantly impact chlamydial growth, the antibiotics data suggest that disruption of ClpP2 via dysregulation of activity is overwhelmingly negative to chlamydial development. However, we cannot exclude a possible effect of these antibiotics on ClpP1 *in vivo*. Clearly, overexpression of ClpP1(S92A) was deleterious to chlamydial growth, thus each ClpP paralog is essential to normal growth and development.

We have shown here an initial characterization of the chlamydial Clp protease system focusing on the ClpP protease components. Overall, the sensitivity of the organisms to perturbations in ClpP activity suggests a critical function for these paralogs in maintenance of chlamydial physiology. We speculate that these systems will also play an integral role in differentiation and possibly persistence (62) (63), which may provide a mechanism that could be leveraged to block growth or eliminate persister cells (46). Further characterization of the chlamydial Clp system could facilitate development of targeted therapeutics for treatment of *C. trachomatis* infection, thereby lowering dependence on broad-spectrum antibiotics. Ongoing efforts are investigating this hypothesis as well as the function of the AAA+ unfoldases.

## Materials and Methods

### Strains and Cell Culture

The human epithelial cell line HEp2 was utilized in the transcriptional and antibiotic studies and was routinely cultivated and passaged in Iscove’s Modified Dulbecco’s Medium (IMDM, Gibco/ThermoFisher; Waltham, MA), and 10% FBS (Sigma; St. Louis, MO). McCoy mouse fibroblasts were used for the purpose of chlamydial transformation, and human epithelial HeLa cells were used for plaque purification of the resulting chlamydial transformants as well as for protein isolation and assessment of ClpP2 expression. All of these cell lines were passaged routinely in Dulbecco’s Modified Eagle’s Medium (DMEM, Gibco/ThermoFisher). Density gradient purified *Chlamydia trachomatis* L2/434/Bu (ATCC VR902B) EBs were used for the antibiotic studies. *C. trachomatis* serovar L2 EBs (25667R) naturally lacking the endogenous plasmid were prepared and used for transformation [see (64)].

### Transcript Analysis Using RT-qPCR

HEp2 cells seeded in 6-well plates were infected at a multiplicity of infection of 1 with *C. trachomatis* serovar L2. At the indicated times post-infection, total RNA and DNA were collected from duplicate wells (63). Briefly, total RNA was collected from infected cells using Trizol reagent, extracted with chloroform, and the aqueous phase precipitated with an equal volume of isopropanol according to the manufacturer’s instructions (ThermoFisher). DNA was removed from total RNA by rigorous DNase-free treatment (ThermoFisher) before 1 μg was reverse transcribed with Superscript III RT (ThermoFisher). Equal volumes of cDNA were used for qPCR. Total DNA was collected from infected cells by trypsinizing the cells, pelleting for 5 min at 400 xg, and resuspending in PBS. Samples were freeze-thawed three times before processing with the DNeasy Blood and Tissue kit according to the manufacturer’s instructions (Qiagen). 150 ng of total DNA were used for qPCR. Transcripts and genomic DNA were quantified by qPCR in 25 μL reactions using 2x SYBR Green Master Mix in an ABI7300 thermal cycler in comparison to a standard curve generated from purified *C. trachomatis* L2 genomic DNA. Transcripts were normalized to genomic DNA.

### C. trachomatis *propagation and detection of Clp proteins*

DFCT28, a GFP-expressing *C. trachomatis* 434/Bu clone (65), was routinely grown in and titered (using the IFU assay) on HeLa cells (66). Briefly, cells were maintained in Dulbecco’s modified Eagle medium (DMEM) supplemented with 10% fetal bovine serum (FBS) and grown at 37 °C with 5% CO_2_. For chlamydial infection experiments, HeLa cells were grown until confluent in 6-well tissue culture dishes and then infected with DFCT28 at an MOI of ~3 using centrifugation at 545 xg for one hour. The infected cells were then incubated at 37 °C with 5% CO_2_ with DMEM/FBS supplemented with 0.2 μg/mL cycloheximide and 1x non-essential amino acids. At various times post infection, the medium was removed, cells were washed twice with 2 ml of PBS, and cells were lysed via addition of 200 μL Laemmli buffer with β-mercaptoethanol followed by heating at 90-100 °C for 5 minutes. Chlamydial protein samples or purified, recombinant ClpP samples were run on 12% SDS-PAGE gels and either stained for total protein with Coomassie Brilliant Blue or transferred to nitrocellulose for western blotting. Blots were probed with a rabbit polyclonal anti-Pa_ClpP1 (*Pseudomonas aeruginosa* designation, similar to ClpP2 from *C. trachomatis*) diluted 1:10000 in 5% milk Tris-buffered saline (mTBS) or anti-Pa_Clp2 (*Pseudomonas aeruginosa* designation, similar to ClpP1 from *C. trachomatis*) at 1:2500. The antibodies were kindly provided by Dr. T. Baker (Massachusetts Institute of Technology) (26). Mouse polyclonal antibodies against chlamydial ClpP1, ClpC, and ClpX (kind gift of Dr. G. Zhong, University of Texas Health Sciences Center at San Antonio) were diluted 1:2000 in mTBS. After incubating with primary antibodies, blots were washed with Tween (0.5%)-TBS (TTBS) and then probed with a goat, anti-rabbit IgG poly-HRP conjugated secondary antibody (Thermo Scientific 32260) diluted 1:1000 in mTBS or goat anti-mouse IgG HRP conjugated secondary antibody (Millipore AP124P) diluted 1:2000 in mTBS. As a control for chlamydial protein, blots were also probed for the major outer membrane protein (MOMP) using a mouse monoclonal anti-MOMP antibody (1:1000; Abcam, ab41193) and a goat anti-mouse IgG HRP conjugated secondary antibody as above. After incubation with the secondary antibodies, blots were washed with TTBS followed by TBS and then incubated with chemiluminescent substrate (EMD Millipore Immobilon ECL) for imaging on a Bio-Rad Chemidoc MP.

### Bioinformatics Analysis

Gene maps and sequences of genes of *Chlamydia trachomatis* used were obtained from STDGen database (http://stdgen.northwestern.edu). RefSeq protein sequences from *E. coli, B. subtilis, M. tuberculosis,* and *P. aeruginosa* were acquired from the NCBI protein database (https://www.ncbi.nlm.nih.gov/guide/proteins/). The ClpP1 vs. ClpP2 protein alignment to find sequence identity was performed using NCBI Protein BLAST function (https://blast.ncbi.nlm.nih.gov/Blast.cgi) (67). Multiple sequence alignments were performed using Clustal Omega (68) with default settings and were presented using Jalview Version 2 (69). PDB files for predicted 3D structures were acquired from the Phyre2 website (http://www.sbg.bio.ic.ac.uk/phyre2/html/page.cgi?id=index) (70). Protein models and model alignments were rendered using the UCSF Chimera package from the Computer Graphics Laboratory, University of California, San Francisco (supported by NIH P41 RR-01081) (71).

### Plasmid Construction

A full list of the primers and plasmids used is included in the supplementary material (S1). Plasmids for the Bacterial Adenylate Cyclase Two-Hybrid (BACTH) system were cloned using the Gateway^®^ recombination system (72). The genes were amplified from *Chlamydia trachomatis* L2 genomic DNA with primers designed to add an *attB* recombination site on either side of the gene. The PCR products were then incubated with a pDONR^TM^221 entry vector (containing *attP* recombination sites) in the presence of BP Clonase II (Invitrogen) that inserts the gene via the flanking *attB* sites and removes *ccdB* endotoxin flanked by the *attP* sites encoded on the plasmid, allowing for positive selection. The result of the BP reaction was an entry vector containing the gene of interest flanked by *attL* sites. 2 μL were transformed into DH5α chemically competent *E. coli* and plated onto an LB agar plate containing 50 μg/mL kanamycin. Plasmid from an individual colony was purified and used for the LR reaction into one of three destination vectors (pST25-DEST, pSNT25-DEST, or pUT18C-DEST). The same entry vector for any given gene was used for all three LR reactions to insert into the destination vector with LR Clonase II. 150 ng of the entry vector was incubated with 150ng of destination vector for 1 hour at room temperature. 2 μL were used to transform XL1 *E. coli*, which were plated on the appropriate selection plate. Purified plasmid from an individual colony was sequence verified prior to use the in BACTH assay (see below).

Constructs for chlamydial transformation were created using the HiFi Cloning (New England Biolabs) protocol. To add the C-terminal 6xHis tag to the Clp proteins, the genes were amplified from the genome with a primer to add the poly-histidine tag. These products served as the template for PCR reactions to add the necessary overlap for the HiFi reaction. Primers were generated using the NEBuilder^®^ assembly tool available from New England BioLabs (http://nebuilder.neb.com). The backbone used was the pTLR2 derivative of the pASK plasmid (36). The pTLR2 backbone was digested using FastDigest BshTI and Eco52I restriction enzymes and dephosphorylated using FastAP (ThermoFisher), and then 25ng of digested plasmid was incubated with a 2:1 ratio of insert copy number to backbone and 2x HiFi master mix (NEB). Following a 15 minute incubation of the reaction mix at 50° C, 2 μL of the reaction was transformed into DH5α chemically competent *E. coli* (NEB) and plated on the appropriate antibiotic selection plate. Positive clone sequences were verified by Eurofins Genomics. Sequence verified plasmids were transformed into *dam-/dcm-E. coli* (New England BioLabs) in order to produce demethylated plasmid, which was sequence and digest verified prior to transformation into *C. trachomatis* (see below).

Strains created or used in this study are listed in the supplementary material. *E. coli* strains were maintained on LB agar plates and grown in LB medium or on LB agar plates supplemented with 100 μg/ml ampicillin as needed. Chlamydial genomic DNA for cloning was obtained from EBs using a phenol:chloroform extraction after extensive heat treatment in the presence of proteinase K (73), and *E. coli* genomic DNA was isolated using sodium hydroxide lysis of colonies. The chlamydial *clpP1* and *clpP2* along with *clpP* from *E. coli* were amplified via PCR using the primers listed in Supplemental Information and Phusion DNA polymerase (Thermo Scientific). PCR products were cloned into the pLATE31 expression vector from Thermo Scientific as directed by the manufacturer to create fusion proteins with a C-terminal 6x His-tag. Plasmids were initially transformed into *E. coli* NEB10 and selected on LB agar ampicillin plates. Transformants were screened for inserts using colony PCR with Fermentas Master Mix (Thermo Scientific) and positive clones were grown for plasmid isolation (GeneJet Plasmid Miniprep Kit, Thermo Scientific). DNA inserts were sequenced by Macrogen USA and sequence-verified plasmids were then transformed into *E. coli* BL21(DE3) bacteria for protein production.

### Chlamydial Transformation

The protocol followed was a modification of the method developed by Mueller and Fields (74). For transformation, 10^6^ *C. trachomatis* serovar L2 EBs (25667R) naturally lacking the endogenous plasmid were incubated with 2 μg of unmethylated plasmid in a volume of 50 μL CaCl_2_ at room temperature for 30 minutes. Each reaction was sufficient for a confluent monolayer of McCoy cells in one well of a six well plate that had been plated a day prior. The transformants were added to a 1 mL overlay of room temperature HBSS per well, and an additional 1 mL of HBSS was then added to each well. The plate was centrifuged at 400 xg for 15 min at room temperature, where the beginning of this step was recorded as the time of infection. Following the spin, the plate was incubated for 15 minutes at 37° C. This infection was recorded as T_0_. The inoculum was aspirated at the end of the incubation and replaced with antibiotic-free DMEM+10% FBS. 8 hours post-infection, the media was replaced with DMEM containing 1 μg/mL cycloheximide, 10 μg/mL gentamicin, and 1 U/mL penicillin. Cells infected with transformants were passaged every 48 hours until a population of penicillin resistant bacteria was established. These EBs were then harvested and frozen in sucrose/phosphate (2SP; (64)) solution at −80° C prior to titration.

### Determining the Effect of Overexpression of Wild Type and Mutant Clp Proteins via Immunofluorescence and Inclusion Forming Unit Analysis

Transformed *C. trachomatis* containing a mutant *clp* gene under control of an anhydrotetracycline (aTc) inducible promoter was used to infect a monolayer of HEp2 cells on coverslips with penicillin as a selection agent. Samples were induced with increasing, subtoxic amounts of aTc at 10 hours post-infection (hpi) and were methanol fixed after a 14 hour pulse (24 hpi). Fixed cells were incubated with an anti-Ctr serovar L2 guinea pig primary antibody (kindly provided by Dr. Rucks, UNMC) to stain for the organism and a goat anti-guinea pig Alexa488 conjugated secondary antibody for visualization of organisms within the inclusion. Additionally, a mouse anti-6xHis tag was used, followed by a goat anti-mouse Alexa594 secondary antibody for confirmation of protein expression and localization. Finally, the samples were stained with DAPI to visualize the host cell and bacterial DNA. Representative images were taken on a Zeiss LSM 800 laser scanning confocal microscope with a 60x objective and 3x digital zoom and were equally color corrected using Adobe Photoshop CC. To assess the effect of wild type and inactive Clp mutant overexpression via the Inclusion Forming Unit (IFU) assay, HEp2 monolayers were infected as described above. At 10hpi, samples either were or were not induced with a 10 nM aTc concentration. Infections were allowed to proceed for another 14h (24hpi) prior to harvest. To harvest IFUs, sample wells were scraped in 2SP and lysed by vortexing with three 1mm glass beads for 45s. Samples were serially diluted in 2SP and titrated in duplicate directly onto a new monolayer of HEp2 cells. Following a 24h incubation, the samples were fixed and stained with an anti-Ctr serovar L2 guinea pig primary and Alexa488 secondary. 15 fields of view were counted for each duplicate well, giving a total of 30 fields of view per experiment. Three independent replicates were performed, and the totals for each experiment were averaged. Values were expressed as a percentage of the uninduced sample to provide an internal control. A Student’s two-tailed t test to compare the induced samples to the uninduced control was performed using the averages of each biological replicate. An asterisk (^*^) indicates P<0.05, N.S. indicates P>0.05.

### Determining Protein-Protein Interactions with the BACTH System

To test interaction of the Clp proteins, the Bacterial Adenylate Cyclase Two-Hybrid (BACTH) assay was utilized. This assay relies on reconstitution of adenylate cyclase activity in adenylate cyclase deficient (Δ*cya*) DHT1 *E. coli*. The genes of interest are translationally fused to one of either subunit, denoted as T18 and T25, of the *B. pertussis* adenylate cyclase toxin. Each wild-type or mutant *clpP* gene cloned into one of the pST25, pSNT25, or pUT18C Gateway^®^ vectors was tested for both homotypic and heterotypic interactions (10). One plasmid from the T25 background and one from the T18 background were co-transformed into the DHT1 CaCl_2_ competent *E. coli* and were plated on a double antibiotic minimal M63 medium selection plate supplemented with 0.5 mM IPTG for induction of the protein, 40 μg/mL Xgal to give a visual readout upon cleavage, and 0.2% maltose as a unique carbon source. These plates also contain 0.04% casein hydrolysate to supplement the bacteria with the branched chain amino acids as Δ*cya* DHT1 *E. coli* cannot synthesize these amino acids. Blue colonies were indicative of positive interaction between proteins since both the *lac* and *mal* operons require reconstituted cAMP production from interacting T25 and T18 fragments to be expressed. Leucine zipper motifs were used for controls in pKT25 and pUT18C backgrounds on the appropriate antibiotic selection plates because these have been previously shown to interact (75). β-galactosidase activity was then measured from 8 colonies to quantify protein interactions. Random positive colonies were used to inoculate individual wells and grown 24 hours at 30° in M63 with 0.2% maltose and appropriate antibiotics. These bacteria were permeabilized with 0.1% SDS and chloroform prior to addition of 0.1% o-nitrophenol-β-galactoside (ONPG). The reaction was stopped using 1 M NaHCO_3_ after precisely 20 minutes of incubation at room temperature. Absorbance at the 405 wavelength was recorded and normalized to bacterial growth (OD_600_), dilution factor, and time (in minutes) of incubation prior to stopping the reaction. Totals were reported in relative units (RU) of β-galactosidase activity.

### Purification of Recombinant wild-type and mutant ClpP proteins

His-tagged Ctr ClpP1, Ctr ClpP1(S92A), Ctr ClpP1(M6L/M7I), Ctr ClpP2, Ctr ClpP2 (S98A), and Ec ClpP were purified from 500 mL cultures of BL21(DE3) *E. coli* transformed with the respective plasmid based on the protocol described in (76). Samples were induced with 0.5 mM IPTG and incubated with shaking for 20 hours at 18°C. Cultures were pelleted and frozen at −20°C prior to purifications. Buffers used are listed in Table S2. Samples were suspended, sonicated, bound to HisPur Cobalt Resin (Thermo Scientific), and washed in buffer A. Proteins were eluted from the resin using buffer B. Buffer exchange for buffer C was performed using a Millipore Amicon Ultra 15 filtration units (3 kDa cut-off). ClpP proteins were quantified using the Bio-Rad Protein assay, assessed for purity on 12% SDS-PAGE gels with Coomassie staining, and identified using anti-His-tag western blot. Blotting was performed using a mouse monoclonal anti-6x His antibody (1:1000; Millipore HIS.H8) and a goat anti-mouse IgG HRP conjugated secondary antibody (1:2000). Protein samples were aliquoted and stored at −80°C.

### In Vitro *Analysis of ClpP Homo-Oligomerization*

5 μg of purified protein was incubated in buffer D at 37°C for 1 hour before being mixed with a 5x native sample buffer (5 mM Tris [pH 6.8], 38 mM glycine, 0.06% bromophenol blue) and analyzed on a BioRad MiniProtean 4-20% gradient gel for Native-PAGE. Samples were run for 90 minutes at 200V. Gels were assessed using Coomassie staining.

### *Assessment of ClpP activity* in vitro

Fluorometric peptide assay: The ClpPs (at 1 μM monomeric concentration) were added to 500 μM of Suc-Luc-Tyr-AMC (Boston Biochem) dissolved in buffer E (50 mM Tris-HCl [pH 8], 200 mM KCl, and 1 mM DTT) or buffer F (with 0.2 M sodium citrate) (44). Final reaction volumes were 50 μl. Reactions were monitored over six hours at 37 °C using a BioTek Synergy HT plate reader set at an excitation of 340/360 and an emission of 440/460 with readings taken at five-minute intervals. Casein degradation assays: Casein (Sigma-Aldrich) was dissolved in buffer E and 1 μg was used per assay. Samples containing casein and 1 μM of the respective ClpP monomer were incubated at 37 °C for 3 hours with or without the respective ACP compound (500 μM). Reactions were halted by mixing with 2x Laemmli buffer containing β-mercaptoethanol and heating at 90-100 °C for 5 minutes. Samples were analyzed for digestion of casein using 12% SDS-PAGE gels followed by Coomassie staining.

### Effect of ACP compounds on chlamydial growth and host cell viability

Antibiotic stocks of ACP1, ACP1a, and ACP1b were synthesized as described (47), resuspended at 25 mg/mL in DMSO, and frozen at −20°C in aliquots to avoid freeze-thawing. Methods for the synthesis, purification, and analysis of these compounds is available in Supplementary Information. For assessment of cell viability upon treatment, four wells of a 96-well plate with a confluent monolayer of HEp2 cells were either infected with density gradient purified wild type *C. trachomatis* serovar L2 with an MOI of 1 or left uninfected. These wells were either treated or not with 25 or 50 μg/mL of ACP1, ACP1a, or ACP1b, with a set of DMSO only samples used as a control. Antibiotics were added eight hours post-infection (hpi). At 24 hpi, 100 μL of 2x Resazurin (Abcam) was added to three wells of each treatment condition, adding only DMEM to the fourth as a background control. Following a four-hour incubation at 37° C, absorbance at the 570 nm wavelength was recorded using a Tecan plate reader. The wells were averaged, subtracting background absorbance from samples without dye. Absorbance was reported as percentage of the untreated samples. To quantify the effect of the drug on *C. trachomatis*, variable treatments of each drug (25 μg and 50 μg) were added or not eight hours post-infection. Cell lysates containing *C. trachomatis* were collected in 2SP and frozen at −80° C prior to serial titration on a fresh cell layer. Inclusion forming units (IFUs), a proxy for recoverable EBs from the initial infection samples, were calculated from the average number of inclusions in 15 fields of view multiplied by the number of fields of view within the well and corrected for the dilution factor and volume of inoculum.

## Acknowledgements

We would like to thank Dr. Elizabeth Rucks of UNMC for providing us with antibodies for analysis by immunofluorescence and for thoughtful review of the manuscript. We thank Dr. Walid Houry (Univ. of Toronto) for providing us with an initial sample of the ACP1b antibiotic, Dr. Tania Baker (MIT) for providing us with Pa_ClpP specific antibodies for use in our experiments, Dr. Gaungming Zhong (UT San Antonio), and Dr. H. Caldwell (NIH/NIAID) for eukaryotic cell lines. Funding for this project was supported by an NSF CAREER award (1810599) to SPO and University funds to DJF. This project was also funded by the Department of Defense office of the Congressionally Directed Medical Research Programs (CDMRP), 2017 Peer Reviewed Medical Research Program (PRMRP) Discovery Award (PR172445) to MCS. Support for the UNMC Advanced Microscopy Core Facility was provided by the Nebraska Research Initiative, the Fred and Pamela Buffett Cancer Center Support Grant (P30CA036727) and an Institutional Development Award (IDeA) from the NIGMS/NIH (P30GM106397).

